# Enhanced disease susceptible variant identification via short identity by descent segments

**DOI:** 10.1101/2023.09.26.559464

**Authors:** Chonghao Wang, Werner Pieter Veldsman, Lu Zhang

## Abstract

Rare diseases affect millions of individuals worldwide, yet diagnostic yields for them still remain low. Among variant identification approaches, identity by descent (IBD) mapping is used to identify disease susceptible variants originating from a recent common ancestor among affected individuals, but existing IBD detection models struggle to identify these variants in short IBD segments. Here, we introduce SILO, a novel model to detect disease susceptible variants in both short and long IBD segments. SILO employs a two-stage procedure to detect IBD segments. In the first stage, SILO identifies long IBD segments based on common variants. In the second stage, SILO utilizes rare variants to detect short IBD segments using a seed-and-extend algorithm. We evaluated SILO in simulated data and real data from the 1000 Genomes Project. Our results demonstrate that SILO outperforms existing models in detecting disease susceptible variants within short IBD segments, and show comparable performance in longer IBD segments. These findings highlight the potential of SILO to increase diagnostic yields for rare diseases by enhancing the identification of previously overlooked disease susceptible variants in short IBD segments.

## Background

Several hundred million individuals worldwide are estimated to be afflicted with rare diseases, approximately 80% of which have a genetic basis [1, 2, 3]. Most of these rare diseases are caused by genomic variants, ranging from single nucleotide variants (SNVs) to structural variations (SVs). Such variants have the potential to affect coding and non-coding regions, leading to dysfunctions in regulatory elements or the formation of aberrant protein products, etc. The advent of next-generation sequencing (NGS) has revolutionized the field of rare disease diagnostics, enabling the identification of numerous disease susceptible variants in previously undiagnosed conditions [4, 5, 6, 7]. In particular, whole-exome sequencing (WES) and whole-genome sequencing (WGS) have proven to be powerful tools in elucidating the genetic basis of rare diseases.

Millions of genomic variants can be identified through NGS, but usually a single variant or a small subset are related to rare disease onset. An naive approach would involve exhaustive wet-lab experimentation to investigate all identified variants; such an approach, however, is infeasible due to cost and technological constraints. Consequently, researchers employ strategies to narrow down the search space and focus on variants that are likely to be associated with the disease of interest. Three primary methods are used: genome-wide association studies (GWAS), linkage analysis, and identity by descent (IBD) mapping. GWAS identifies the genomic variants that are significantly more prevalent in affected individuals compared to control individuals. However, GWAS generally targets common variants (minor allele frequencies (MAFs) >= 5%) that are usually associated with common complex diseases rather than rare diseases. Linkage analysis relies on the co-segregation of genomic variants with the disease in the families. IBD mapping identifies genomic regions that are inherited from an affected common ancestor (i.e. the IBD segments) and are shared among all individuals exhibiting the disease phenotype. The underlying hypothesis of IBD mapping is that a founder possesses disease susceptible variants to a disease, and the regions containing these variants are subsequently transmitted to affected offspring. Consequently, researchers can locate and identify these variants through detecting the IBD segments shared among affected individuals. Both linkage analysis and IBD mapping can be used to identify disease susceptible variants of rare diseases within families; however, IBD mapping is more effective to identify several disease susceptible variants with modest effects than linkage analysis [8]. Furthermore, IBD mapping can be applied to distant relatives and populations, particularly those with a history of inbreeding.

IBD mapping has been successfully applied to identify disease susceptible variants in various contexts. For example, Krawitz *et al*. [9] employed IBD mapping in siblings affected by hyperphosphatasia mental retardation syndrome and identified disease susceptible variants in *PIGV* gene. Hsueh *et al*. [10] used IBD mapping to discover disease susceptible variants in *APOC3* gene that resulted in serum triglycerides, providing an explanation for the results obtained from previous GWAS. In a research conducted by Belbin *et al*. [11], they detected a disease susceptible variant in *COL27A1* gene that caused a collagen disease in Puerto Ricans through IBD mapping in a biobank dataset. Furthermore, Henden *et al*. [12] used WGS data from 83 cases across 25 families and 3 sporadic cases to identify disease susceptible variants in *SOD1* gene associated with amyotrophic lateral sclerosis. These studies highlight the effectiveness of IBD mapping in locating disease susceptible variants across a range of rare genetic diseases.

Despite the advancements in the field of rare disease diagnostics, the overall diagnostic yield remains below 50% [3, 6, 13, 14]. The low diagnostic yields can be attributed to several factors, among which a notable one is the challenge to identify a few truly disease susceptible variants amidst numerous identified genomic variants. This necessitates a prioritization process where researchers need to determine which variants warrant further analysis. IBD mapping is one of the approaches for prioritization. However, the prioritization process carries the risk of missing disease susceptible variants if they are not included in the prioritized group, resulting in undiagnosed conditions. Existing IBD detection models [15, 16, 17, 18, 19, 20, 21, 22] primarily focus on detecting long IBD segments (> 2 centimorgan (cM)); however, they may lack sufficient power to identify shorter IBD segments [20]. As a result, if a disease susceptible variant is located within a short IBD segment between two affected individuals, it can be dismissed if the short IBD segment remains undetected.

The lengths of IBD segments inherited from a common ancestor can be approximated by an exponential distribution, with a mean length of 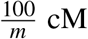, where *m* stands for the number of meioses that have occurred since the most recent common ancestor. Consequently, individuals with a common ancestor 25 generations (i.e. 50 meioses) ago are expected to share an IBD segment with an average length of 2 cM, while Browning and Browning [23] noted that fifth cousins often lack detectable IBD segments in practice. Historically, there has been a lack of interest in short IBD segments as they were thought to be relics of distant ancestry and were relatively rare. Nonetheless, our simulation study suggested that short IBD segments accounted for 10% to 20% of the total IBD segments shared between second cousins to sixth cousins, with approximately 70% of these segments containing at least one rare variants (MAF < 1%) where the minor alleles were shared (Table 1). In addition, high genotype error rates in certain regions can potentially fragment long IBD segments into smaller pieces that may go undetected. Overlooking these short IBD segments can dismiss the potential disease susceptible variants they contain, thereby leaving some rare diseases undiagnosed.

**Table 1:** Averaged proportions of short IBD segments and rare variants within these segments across various degrees of kinship, calculated from 100 simulated pedigrees.

| Relationship | Short IBD segments proportion | Rare variants proportion |
| --- | --- | --- |
| Full sibling | 0.029 | 0.681 |
| First cousin | 0.066 | 0.681 |
| Second cousin | 0.105 | 0.698 |
| Third cousin | 0.141 | 0.703 |
| Fourth cousin | 0.171 | 0.760 |
| Fifth cousin | 0.220 | 0.649 |
| Sixth cousin | 0.160 | 0.769 |
| Average | 0.127 | 0.706 |

In response to this problem, we implemented SILO, a novel algorithm to detect short IBD segments that contain rare variants, as well as long IBD segments. SILO employs an approach similar to that used by Parente2 to identify long IBD segments (e.g. lengths >= 2 cM) using common variants, but adapts the approach to detect short IBD segments using rare variants (**Methods**). SILO assumes that the probability of the presence of a minor allele of a rare variant follows a Bernoulli distribution, with its parameter modeled by a Beta distribution. SILO implements a seed-and-extend algorithm to identify short IBD segments and utilizes a cleaning procedure to remove false positive IBD segments. By leveraging the IBD segments inferred from SILO among affected individuals, the potential disease susceptible variants residing in both short and long IBD segments can be identified. To investigate the ability on finding disease susceptible variants, we compared SILO with four state-of-the-art IBD detection models (hap-IBD [20], GERMLINE2 [18], TRUFFLE [22], and Parente [15]) on simulated data and on a real pedigree from the 1000 Genomes Project. SILO demonstrated a remarkably superior performance in detecting disease susceptible variants within short IBD segments compared to the other models. It achieved a mean hit rate that was 8.3 times higher than the second-best model using default hyperparameters for IBD lengths between 1 to 2 cM in simulated pedigrees. In the exercise on a real three-generation pedigree from the 1000 Genomes Project, we demonstrated that SILO identified nine rare variants, among which one variant was predicted to be pathogenic by CADD v1.7 [24], that was overlooked by all other models. In contrast, the other models did not detect any potential disease susceptible variants missed by SILO. These results underscore SILO’s unparalleled ability to detect disease susceptible variants in short IBD segments. By identifying these elusive variants, SILO has the potential to improve diagnostic yields for rare diseases.

## Results

### Overview of SILO

SILO employs a two-stage procedure to identify both long and short IBD segments between a pair of individuals. In the first stage, SILO identifies long IBD segments based on common variants. The common variants in one chromosome are divided into contiguous windows (without overlap) with the same number of common variants. SILO uses inner log-likelihood ratios (inner-LLRs) to determine the likelihood of a window being IBD versus non-IBD. Due to high variances of inner-LLRs within certain windows, SILO converts inner-LLRs to outer log-likelihood ratios (outer-LLRs) using empirical distributions. A sliding block (one window for each step) is defined to include several contiguous windows until their total length equals or slightly exceeds the pre-defined block length threshold (3 cM by default). The block outer-LLR is obtained by combining the outer-LLRs from all the included windows. When the block outer-LLR surpasses a predefined threshold value (*T*_*LLR*_), it signifies that this specific block is identified as an IBD segment between the two individuals.

In the second stage, SILO seeks to detect short IBD segments with rare variants using a seed-and-extend algorithm. A seed is defined as a window (i) with shared minor alleles of rare variants between two individuals; (ii) with the window outer-LLR not smaller than a threshold *T*_*w*1_; (iii) with a combined-LLR (the window outer-LLR and rare variant LLRs) not smaller than *T*_*LLR*_. SILO extends from the seed to its surrounding windows and terminates until certain conditions are met (**Determination of IBD segments**). Moreover, SILO employs a cleaning procedure to remove potentially false positive IBD regions based on discordant alleles of rare variants. The workflow of SILO is shown in Figure 1.

**Figure 1:**
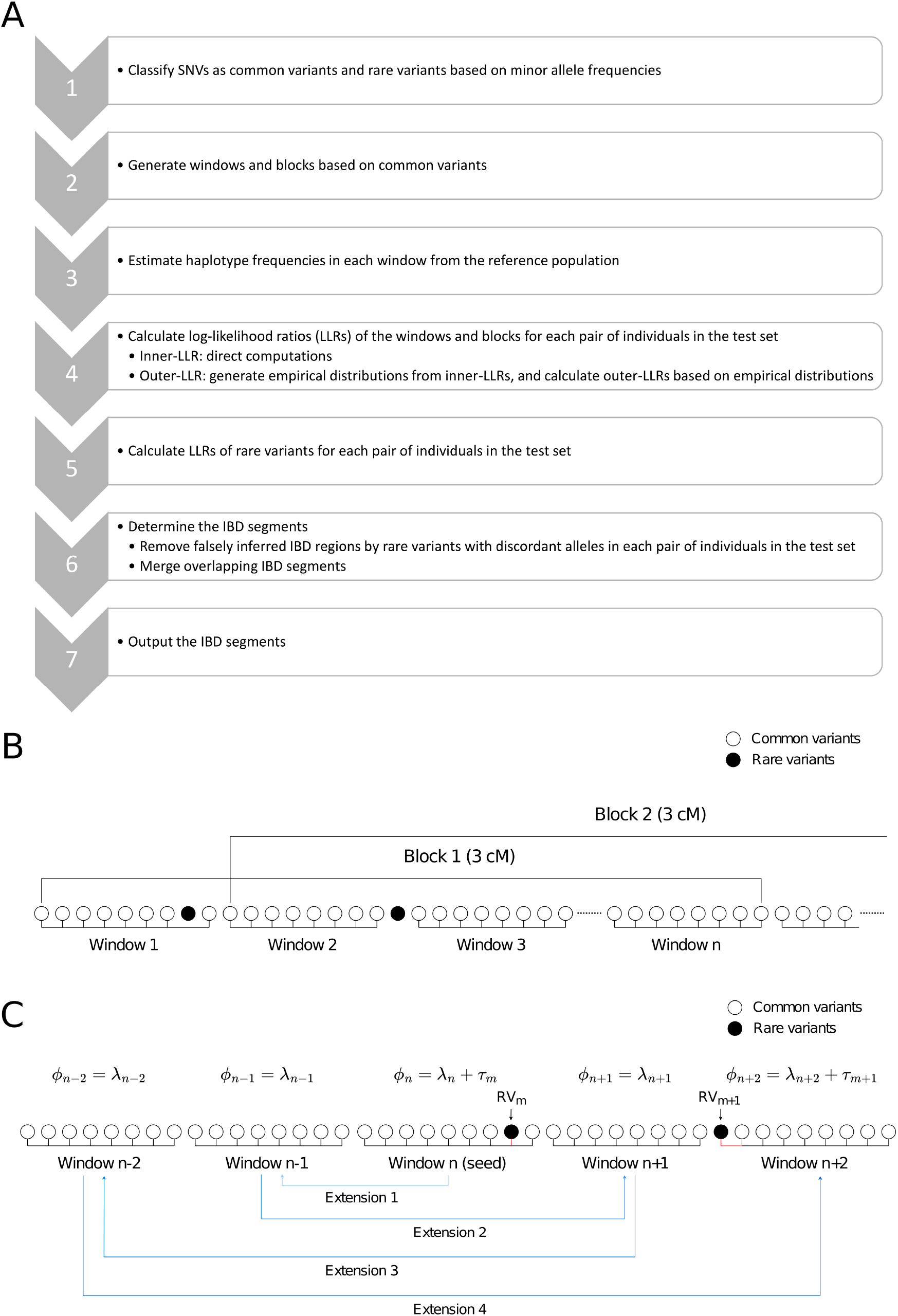
SILO framework. (**A**) The SILO algorithm. (**B**) SILO’s window-and-block structure. A chromosome is divided into many windows. Each window contains a fixed number of common variants (eight in this figure). A block is built on top of windows, and its length is measured in centimorgan (3 cM in this figure). (**C**) Seed-and-extend diagram. During the extension, SILO first extends to the left flanking window and then to the right flanking window. If the extension to a window stops, the extension in that direction stops.

### SILO detects a disease susceptible variant overlooked by existing models in a simulated pedigree

We employed ped-sim to simulate a 4-generation pedigree, inclusive of individuals harboring a disease susceptible variant. The haplotypes of the founders in this pedigree were extracted from white people of the 1000 Genomes Project [25]. The structure of the simulated pedigree is depicted in Figure 2A. As shown in Figure 2B, the disease susceptible variant first emerged in *g1i2* that precipitated the onset of the disease. This disease susceptible variant was subsequently passed down to six affected descendants (*g2i2, g2i3, g3i2, g3i3, g4i1, g4i2*). Notably, a short segment of the original haplotype from *g1i2*, which harbored the disease susceptible variant, was inherited by *g4i2*. Consequently, *g4i2* shares a short IBD segment with all affected individuals within this pedigree. As demonstrated in Figure 2C, for the pair *g4i1* and *g4i2*, only SILO successfully detected the IBD segment containing the disease susceptible variant. GERMLINE2 identified a portion of the IBD segment, but it did not capture the region containing the disease susceptible variant. Parente, TRUFFLE, and hap-IBD all failed to identify any IBD segments. These results reflected that SILO could detect disease susceptible variants in short IBD segments that were missed by the other models, thereby potentially increasing the diagnostic yield for rare diseases.

**Figure 2:**
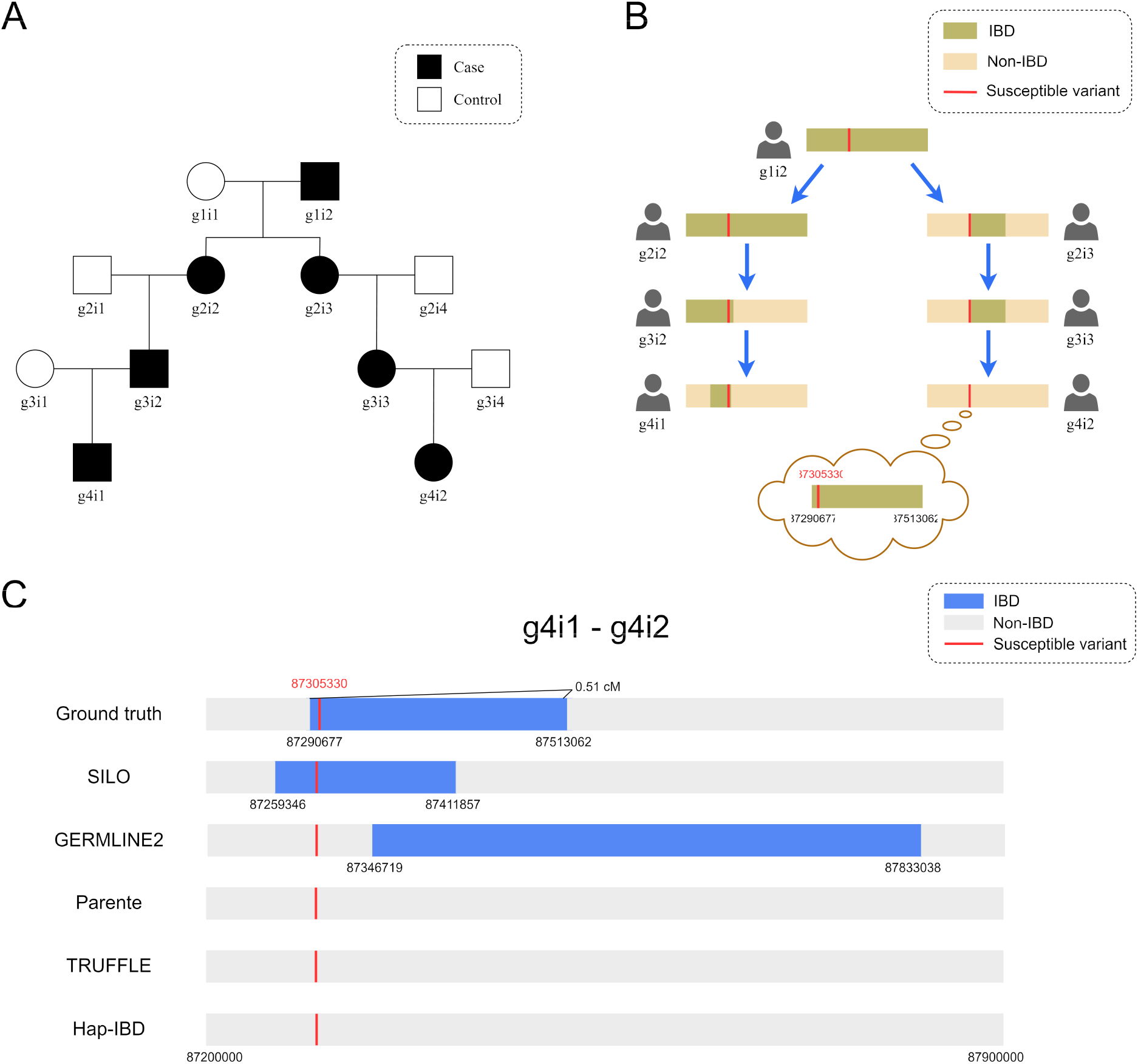
(**A**) The simulated pedigree structure. (**B**) Transmission of a disease susceptible variant. Each IBD segment delineates the inherited region from the founder *g1i2*. (**C**) The ground truth and inferred IBD segments from each model.

### SILO outperforms existing models in simulated pedigrees

In addition to the simulated pedigree described above, we conducted a larger scale evaluation of each model by simulating a series of pedigrees. Specifically, we used ped-sim to simulate 50 pedigrees, where a subset of individuals in each pedigree harbored a disease susceptible variant (randomly selected from rare variants), and at least one pair of individuals within each pedigree shared an IBD segment longer than 2 cM that encompassed the disease susceptible variant. Furthermore, we simulated 25 pedigrees with at least one pair shared an IBD segment between 1 to 2 cM, and 25 pedigrees with IBD segments shorter than 1 cM. As a result, we generated 50 pairs of individuals sharing an IBD segment longer than 2 cM that covered a disease susceptible variant, 25 pairs with IBD segments between 1 to 2 cM, and 25 pairs with IBD segments shorter than 1 cM.

We tested two sets of hyperparameters for GERMLINE2, Parente, TRUFFLE and hap-IBD: the default hyperparameters and a set of tuned hyperparameters (listed in supplementary tables). As demonstrated in Figure 3A, SILO achieved mean hit rates of 1 and significantly outperformed the other models for IBD lengths between 1 to 2 cM and shorter than 1 cM. Only GERMLINE2 and Parente exhibited comparable results to SILO for IBD lengths longer than 2 cM. With default hyperparameters, SILO demonstrated a mean hit rate 8.3 times higher than the second-best model, GERMLINE2, for IBD lengths between 1 to 2 cM, and 3.6 times higher than GERMLINE2 for IBD lengths shorter than 1 cM. With tuned hyperparameters, SILO maintained a mean hit rate nearly 2 times higher than the GERMLINE2 for IBD lengths between 1 to 2 cM, and 1.3 times higher than Parente for IBD lengths shorter than 1 cM. Besides evaluating the ability to detect potential disease susceptible variants, we assessed the proportion of rare variants that SILO accurately excluded from disease susceptibility consideration based on the inferred non-IBD regions. Our results demonstrated that a maximum of more than 25% rare variants with minor alleles shared between a pair of individuals were excluded by SILO.

**Figure 3:**
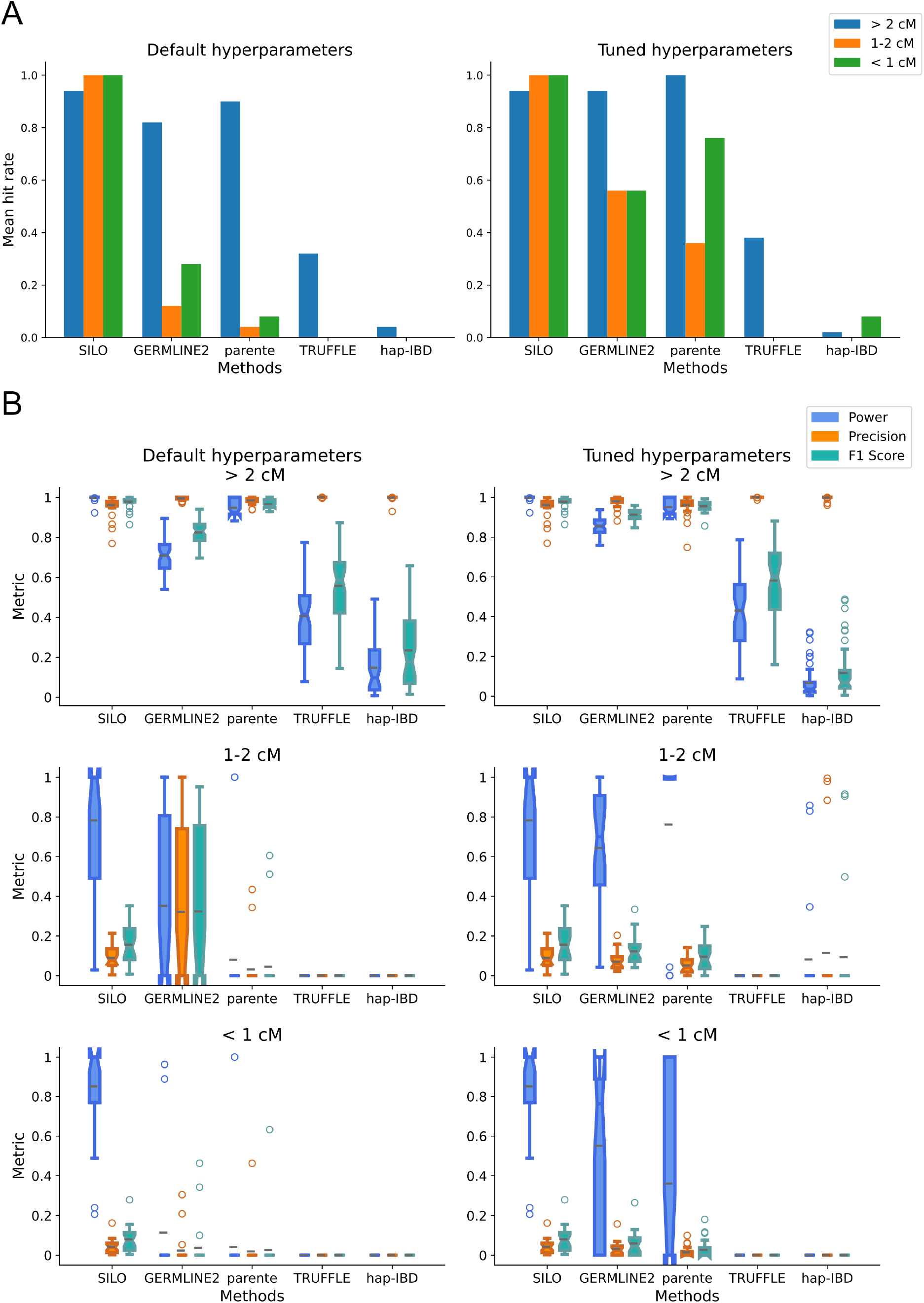
(**A**) The mean hit rates of disease susceptible variants for simulated pedigrees with IBD segments longer than 2 cM, between 1 to 2 cM, and shorter than 1 cM. The performance of IBD detection models with default hyperparameters and tuned hyperparameters are shown in two plots, separately. (**B**) Power, precision, and *F*_1_ scores of each model for the simulated pedigrees. 20

We also evaluated the performance of each model in detecting IBD segments, as measured by power, precision, and *F*_1_ scores. As shown in Figure 3B, SILO achieved a mean power of 0.998, a mean precision of 0.961, and a mean *F*_1_ score of 0.979 for IBD lengths longer than 2 cM, slightly surpassing the second best model, Parente. For IBD lengths between 1 to 2 cM, SILO demonstrated relatively better performance than the other models with tuned hyperparameter setting, exhibiting higher mean power, precision, and *F*_1_ scores. For IBD lengths shorter than 1 cM, GERMLINE2, Parente, TRUFFLE, hap-IBD all struggled to detect any IBD segments with default hyperparameter setting, indicating their potential to overlook disease susceptible variants within such short IBD segments in practice. In contrast, SILO achieved a mean power of 0.85 and exhibited a higher mean precision and *F*_1_ score compared to other IBD detection models. In summary, SILO demonstrated superiority to other top-performing models in identifying disease susceptible variants and IBD segments for IBD lengths shorter than 2 cM, while exhibiting comparable performance for IBD lengths longer than 2 cM.

### SILO achieves the best performance in simulated pairs with no latent IBD segments

To mitigate the effects of latent IBD segments that might be present in the simulated data due to the selection of founders from the 1000 Genomes Project, we employed Fastsimcoal2 [26, 27] to simulate a reference population of 1,000 individuals and a test set of 150 individuals. We removed latent IBD segments of the tested pairs in the test set and then implanted short IBD segments encompassing disease susceptible variants to evaluate the performance in identifying these variants and IBD segments. To investigate the performance of each IBD detection model across different IBD length scales, we selected four ranges: 0.1 cM-0.2 cM, 0.2 cM-0.5 cM, 0.5 cM-1 cM and 1 cM-2 cM. In total, four experiments were conducted, each with 100 replicates. As depicted in Figure 4A, the mean hit rates of SILO were significantly higher than those of the other models for IBD lengths shorter than 1 cM. Notably, SILO achieved an mean hit rate of 0.99 for IBD lengths between 0.1 to 0.2 cM, which was over 8 times higher than the second-best model, hap-IBD. As the IBD lengths increased, the mean hit rates of the other models also improved accordingly. For IBD lengths longer than 1 cM, Parente achieved an mean hit rate of 0.99, which was close to SILO’s hit rate of 1. These results highlighted the superiority of SILO over the other models in identifying disease susceptible variants within short IBD segments.

**Figure 4:**
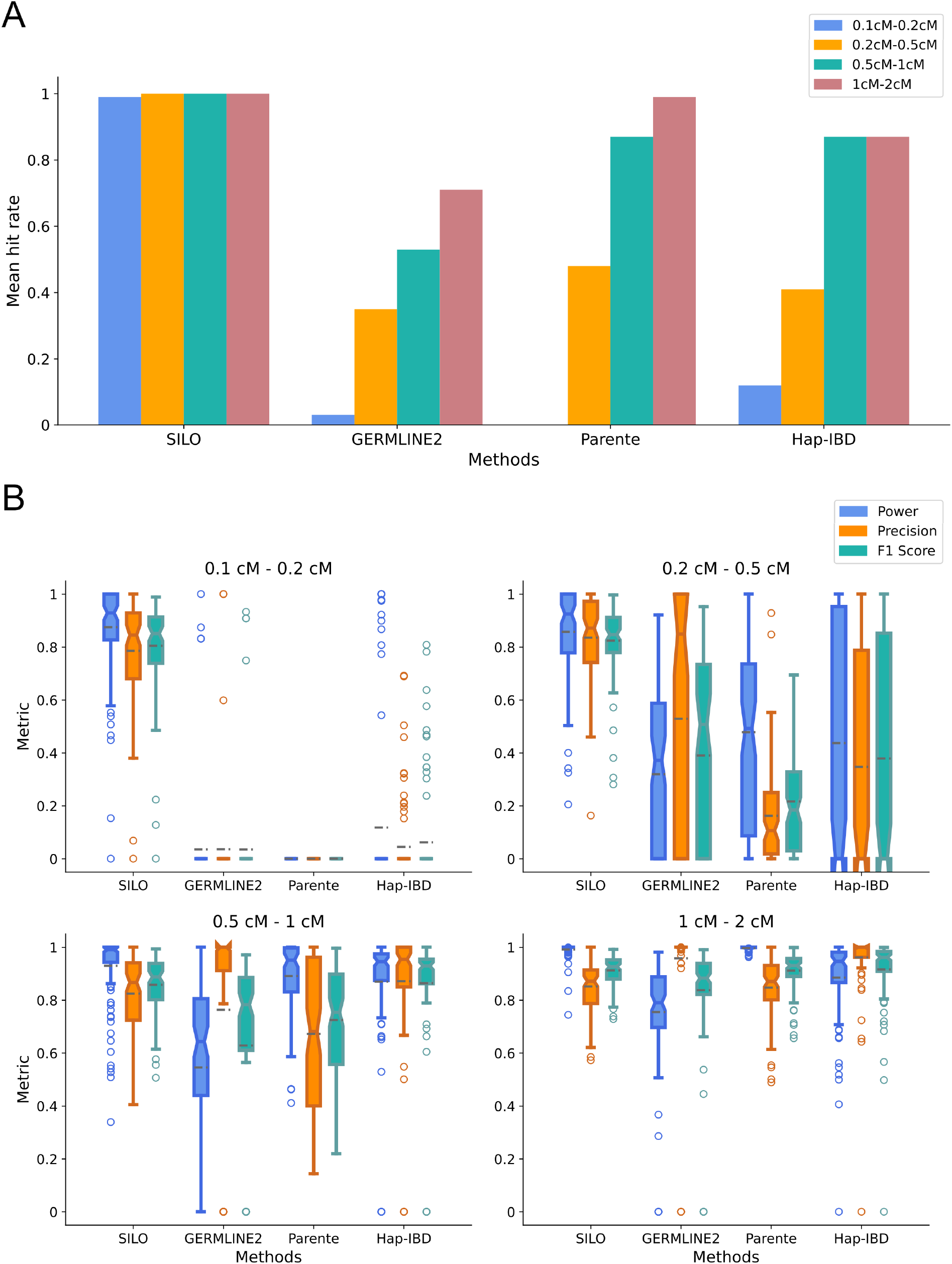
(**A**) The mean hit rates of disease susceptible variants for simulated pairs with no latent IBD segments. Four different lengths of IBD segments encompassing the variants were evaluated: 0.1-0.2 cM, 0.2-0.5 cM, 0.5-1 cM, and 1-2 cM. (**B**) Power, precision, and *F*_1_ scores of each model for the simulated data.

Figure 4B revealed that SILO showed significant improvements over the other models in detecting very short IBD segments. In the experiments targeting the shortest (0.1-0.2 cM) IBD segments, all models underperformed compared to SILO. Hap-IBD, which performed the best among the other models, merely achieved a mean *F*_1_ score of 0.06. In contrast, SILO obtained a mean *F*_1_ score of 0.8. As the length of IBD segments increased, the other models exhibited higher power, precision, and *F*_1_ scores. Despite notable improvements compared to the IBD length of the 0.1 cM-0.2 cM range, the other models still struggled to reliably identify IBD segments between 0.2 to 0.5 cM, where the mean *F*_1_ score of the best performing model was 0.39, while SILO achieved a mean *F*_1_ score in excess of 0.8. When the IBD segment lengths exceed 0.5 cM, some of the other models showed *F*_1_ scores that were comparable to those that were obtained by SILO. Specifically, hap-IBD achieved a mean *F*_1_ score of 0.86 for IBD segments between 0.5 and 1 cM, while hap-IBD, GERMLINE2, and Parente obtained mean *F*_1_ scores of 0.92, 0.84, and 0.91, respectively, for IBD segments between 1 and 2 cM.

Unlike hap-IBD, GERMLINE2 and Parente that achieved at least some inference accuracy, TRUFFLE could not infer any IBD segments in all experiments. It was expected that TRUFFLE would obtain such results as it utilized genotype but not haplotype information and was thus limited to the detection of long IBD segments.

### SILO identifies a disease susceptible variant missed by existing models in real data

In conjunction with the simulation study, we evaluated the performance of SILO on real data. As it was generally challenging to determine the ground truth for IBD segments in real data, we selected six individuals within a Mexican pedigree in the 1000 Genomes Project [25, 28] as the test set. By leveraging the known relationships of these individuals (**Figure S1**), we investigated the transmission of haplotype segments, and evaluated the performance of each model in detecting potential disease susceptible variants missed by the other models.

We ran each model on every pair of the six pedigree members for all autosomes and selected the pair NA19662-NA19686 for analysis. Based on the IBD segments inferred by each model, we investigated the presence of rare variants that could only be identified by one model. To confirm that the rare variants were indeed inherited from one haplotype of a founder, NA19660 or NA19661, we applied the criteria outlined in **Methods**. As a result, SILO detected nine rare variants that were overlooked by the other models, while the other models did not identify any rare variants that were missed by SILO. After evaluating these nine rare variants using CADD v1.7 [24], we identified a potential disease susceptible variant on chromosome 20 that had a PHRED score of 10.97, surpassing the cutoff value of 10 suggested by CADD.

This potential disease susceptible variant was shown in Figure 5. The variant was located at position 162054, a non-coding region, on chromosome 20. SILO inferred a short IBD segment of a length 0.112 cM that covered this variant between NA19662 and NA19686. Multiple SNVs with discordant alleles were present downstream of the segment in a 0.137 cM region, prohibiting the extension of the short IBD segment to a longer one. As only NA19661, NA19662, NA19685, and NA19686 contained a haplotype harboring the minor allele of this variant, we could confirm that NA19662 and NA19686 both inherited this haplotype from NA19661, indicating that this variant was transmitted along with a short IBD segment.

**Figure 5:**
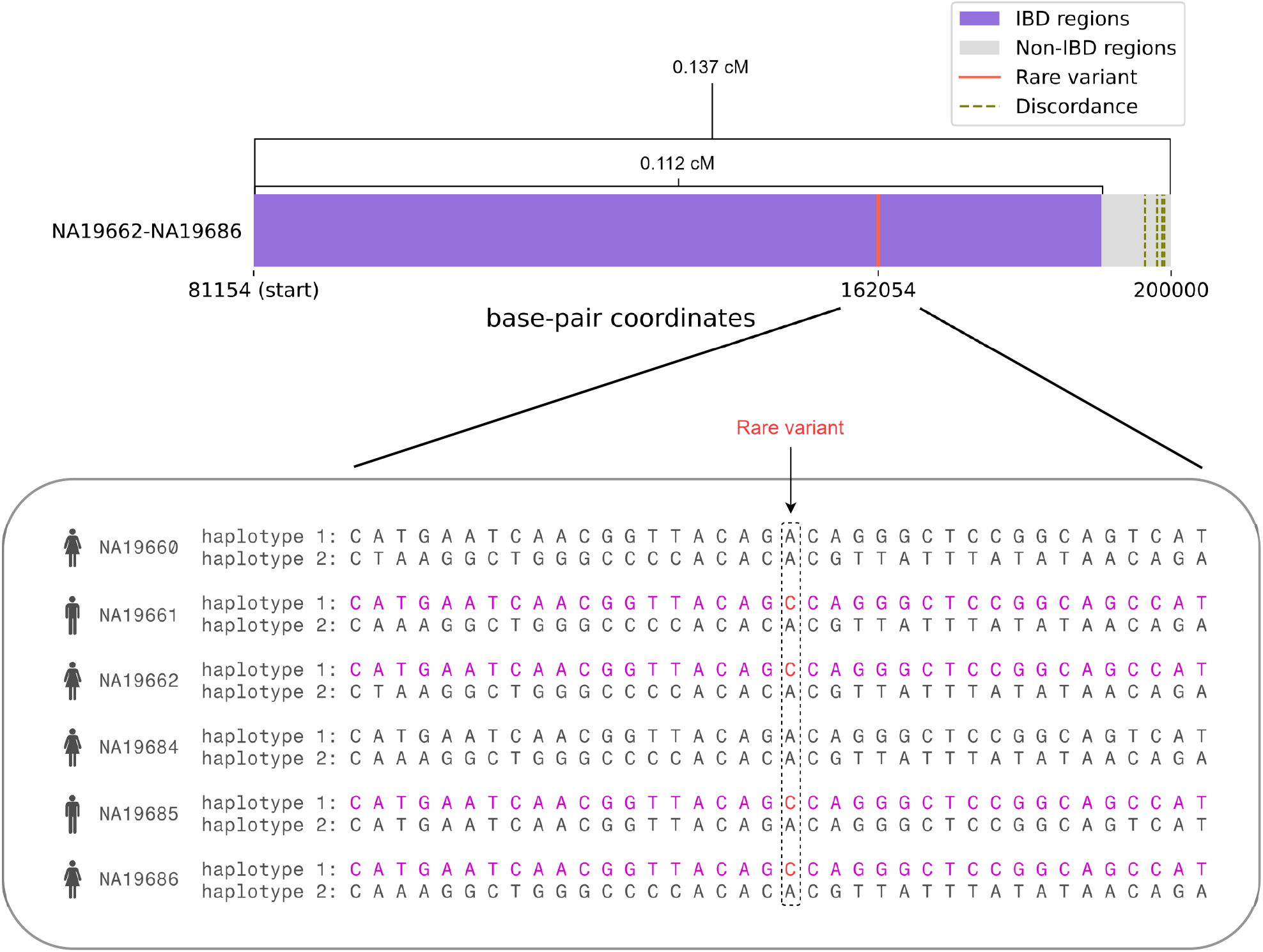
An example of a disease susceptible variant within a short IBD segment inferred by SILO, shared by NA19662 and NA19686 on chromosome 20. This short IBD segment is inherently unextendible due to the presence of multiple SNVs with discordant alleles located downstream.

We also evaluated whether the proportions of IBD segments inferred by each model in the autosomes were generally consistent with the relationships between each pair of the pedigree members. The expected proportion of IBD segments could be derived based on the known relationships of the pedigree members [29]. **Figure S2** showed that the proportions obtained by SILO generally matched the expected proportions. Although there was a noticeable difference between the expected and inferred proportions of IBD segments for NA19660-NA19686 and NA19661-NA19686, the other models showed similar trends to SILO on the proportionality (**Table S1**). For example, the proportions of GERMLINE2 for the two pairs just mentioned were 0.326 and 0.637, respectively, and the proportions of Parente for them were 0.325 and 0.678, respectively. This suggests that either the true proportions for these two pairs differed substantially from their expected proportions or that all models had difficulties in inferring IBD segments for these two pairs.

### The influence of hyperparameters

The performance of SILO is influenced by several hyperparameters. As illustrated in **Figure S3**, variations in the *window size* from 10 to 20 had a modest impact on the performance. In simulated pedigrees, a decrease in power of approximately 0.1 was observed when the *window size* was increased from 14 to 16 for IBD lengths ranging from 1 to 2 cM. Moreover, *window sizes* larger than 16 exhibited inferior performance compared to their smaller counterparts. In addition, incrementing window size improved precision towards an optimal *window size* (around 14) beyond which a decrease in precision was observed.

The thresholds *T*_*w*1_ and *T*_*w*2_, which serve as cutoff values for the differentiation of identity by state (IBS) windows, were found to have large effects on the power of detecting short IBD segment. As presented in **Table S2**, decreasing the value of *T*_*w*1_ from 0 to -0.3 resulted in an increase in power from 0.737 to 0.879 for IBD lengths below 1 cM. Similarly, decreasing *T*_*w*2_ from -0.2 to -0.4 led to an increase in power from 0.793 to 0.880. Comparable trends were also observed for IBD lengths between 1 and 2 cM, as depicted in **Table S3**. However, it is important to note that the reduction in *T*_*w*1_ and *T*_*w*2_ values also corresponded to a decrease in precision. Consequently, a balance must be achieved when selecting the values for *T*_*w*1_ and *T*_*w*2_ to optimize the performance.

## Discussion

The identification of causal variants for rare genetic diseases has long been a significant challenge in clinical practice. Despite advancements in the sequencing technologies and bioinformtics algorithms, the diagnostic yields for the rare diseases have generally been below 50% [3, 6, 13, 14]. As highlighted by AlAbdi et al. [14], many causal variants were missed in the interpretation stage rather than the detection stage, indicating that these variants could be detected by the current technology but were not prioritized or overlooked during the evaluation process. This highlights the need for more sophisticated and comprehensive approaches to identify and prioritize disease susceptible variants.

In the context of IBD mapping, existing IBD detection models primarily focused on identifying long IBD segments (e.g. > 2 cM) and exhibited poor performance in identifying shorter IBD segments. However, our simulation experiments demonstrated the importance of these short IBD segments. We discovered that short IBD segments were indeed present even in pedigrees, accounting for 10% to 20% of total IBD segments across various degrees of kinship. Notably, approximately 70% of these short IBD segments contained rare variants with two individuals sharing minor alleles. When affected individuals share the minor alleles of these rare variants, they are considered potential disease susceptible variants, and causal variants may lie within these variants. The disease susceptible variants in short IBD segments can be easily overlooked by existing models. To address these problems, we present SILO to detect disease susceptible variants in both short and long IBD segments, thereby enhancing the identification of disease susceptible variants.

Genotype errors play a crucial role in the performance of IBD detection models. SILO intrinsically considers the impacts of genotype errors. For common variants, a genotype error rate of 0.5% is sufficient for SILO to make accurate predictions. However, when dealing with rare variants, the presence of high genotype error rates poses a challenge. In such cases, SILO assigns smaller weights to the observed genotypes and increases the weights on other possible genotypes. We investigated the impacts of different genotype error rates on SILO. Our findings revealed that with a genotype error rate of 1%, SILO did not fully utilize the information from rare variants. This can be attributed to the higher likelihood that the sharing of minor alleles of rare variants was due to genotype errors rather than true genetic sharing. Based on our empirical results, a genotype error rate of *ε* enables SILO to extract sufficient information from rare variants with MAFs above 10 *×ε*^2^. For instance, it was observed that a genotype error rate of 0.1% allowed SILO to extract adequate information from rare variants with MAFs above 10^*−*5^, rather than regarding them as genotype errors.

A challenging problem in IBD segment detection is the determination of its boundary points. When SILO detects long IBD segments, the minimum length of an IBD segment is the block length. As a block consists of several windows, it is possible that the windows near the block boundary are non-IBD (e.g. IBS segments). However, if the summation of outer-LLRs of all windows in the block exceeds the threshold, the entire block is considered an IBD segment. For short IBD segments detection, the seed-and-extend algorithm may also include some non-IBD regions. To mitigate this issue, SILO employs a cleaning procedure. This procedure utilizes rare variants with discordant alleles to identify non-IBD regions surrounding them. If inferred IBD segments overlap with these non-IBD regions, the overlapping parts are considered false positives and are removed from the inferred IBD segments.

SILO makes a significant contribution to the field by enhancing the identification of disease susceptible variants in short IBD segments. Our results demonstrated that SILO outperformed state-of-the-art models in detecting disease susceptible variants in short IBD segments, and it achieved comparable performance with these models when detecting the variants in longer IBD segments. In practice, SILO is recommended to be employed in small populations or pedigrees. Based on the IBD segments inferred by SILO among affected individuals, the disease susceptible variants within these segments can be prioritized for further analysis, such as variant effect prediction and multiplexed assays of variant effect. By identifying the previously overlooked variants in short IBD segments, the diagnostic yields for rare diseases can be potentially increased.

## Methods

### Calculation of the outer log-likelihood ratios of common variants

SILO detects long IBD segments based on common variants (MAF>=5%). As illustrated in Figure 1, a chromosome is split into contiguous windows (without overlap) and each window contains *n* common variants. A sliding block (one window per step) is built on top of the windows, and its length is measured in centimorgan. The windows are incorporated in a block until the block length equals or slightly exceeds 3 cM. SILO refers to the similar framework of Parente2 to identify long IBD segments, while it modifies the way to split bins in empirical distributions and removes the augmented windows and window subsets in Parente2 to allow the extension of windows with rare variants for short IBD segment detection.

The probabilities of observing *g*(*w*) and *g*^*′*^(*w*) from IBD and non-IBD segments are defined as

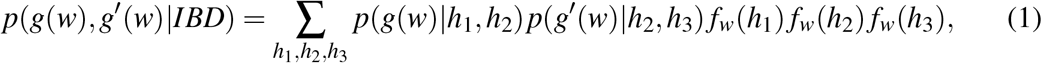

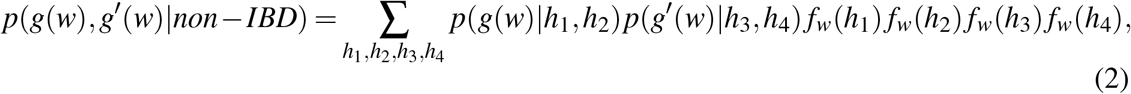

where *p*(*g*(*w*)|*h*_1_, *h*_2_) is defined as

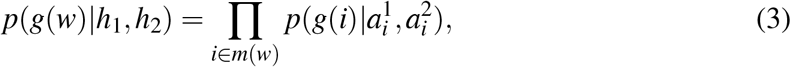

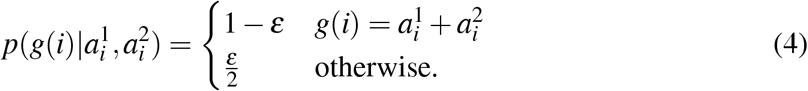

We denote *g*(*w*) and *g*^*′*^(*w*) as the genotypes of common variants in window *w* of two individuals. The genotype of each common variant is represented as 0, 1 and 2 based on the number of major alleles. The haplotypes of two individuals can be denoted as (*h*_1_, *h*_2_) and (*h*_2_, *h*_3_), if they inherited *h*_2_ from the same ancestor; otherwise, they are represented as (*h*_1_, *h*_2_) and (*h*_3_, *h*_4_). *m*(*w*) is a set of all common variants included in window *w*, 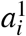 and 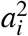 represent the alleles of the common variant *i* from *h*_1_ and *h*_2_. *f*_*w*_(*h*_*i*_) (*i ∈ {*1, 2, 3, 4 *}*) is the frequency of *h*_*i*_ in window *w*, which is estimated in the available haplotypes from the same population (taken as the reference population). *ε* is the genotype error rate of common variants (default *ε* = 0.5%). *p*(*g*^*′*^(*w*) | *h*_2_, *h*_3_) and *p*(*g*^*′*^(*w*) | *h*_3_, *h*_4_) can be defined in the same way.

The inner-LLR (*γ*_*w*_(*g*(*w*), *g*^*′*^(*w*))) of the window *w* is used to compare the magnitude of *p*(*g*(*w*), *g*^*′*^(*w*) |*IBD*) and *p*(*g*(*w*), *g*^*′*^(*w*) | *non −IBD*) and show the degree of differences between them, and it is defined as

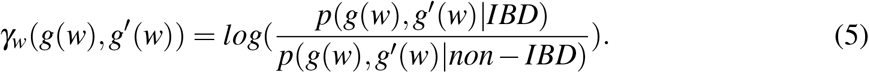

Because some windows in a block may have high variances for *γ*_*w*_(*g*(*w*), *g*^*′*^(*w*)), making the inner-LLR of the block sensitive to such factor, we generate two empirical distributions (*Q*_*I*_ and *Q*_*Ī*_) for *γ*_*w*_(*g*(*w*), *g*^*′*^(*w*)) under the assumption of IBD and non-IBD segments, respectively. *Q*_*I*_ and *Q*_*Ī*_ are generated by two groups of *γ*_*w*_(*g*(*w*), *g*^*′*^(*w*)) values for IBD and non-IBD segments. We assign these *γ*_*w*_(*g*(*w*), *g*^*′*^(*w*)) values to non-overlapping bins to approximate the probability density function of *Q*_*I*_ and *Q*_*Ī*_. We consider the values in (*−* inf, -18) as *bin*_*L*_ and in (18, inf) as *bin*_*H*_ for extreme values, and the remaining bins (bin size=2) span over [-18, 18]. We define outer-LLR (*λ*_*w*_(*g*(*w*), *g*^*′*^(*w*)) as

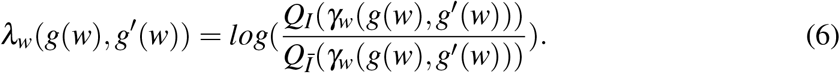

The inner-LLR (*γ*_*b*_) and outer-LLR (*λ*_*b*_) of a block are calculated by summing up the corresponding value for each involved window, based on the independence assumption among these windows.

### Calculation of the log-likelihood ratios of rare variants

We assume rare variants are in trivial LD with the other variants. Therefore, we calculate probabilities and LLR for individual rare variants instead of windows. The probabilities of observing *g*(*m*) and *g*^*′*^(*m*) from IBD and non-IBD segments are

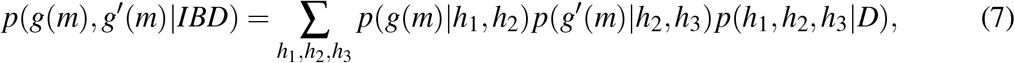

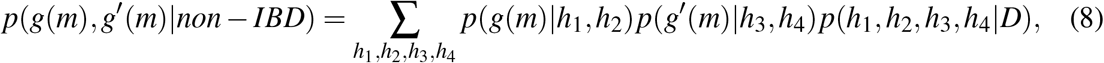

where *g*(*m*) and *g*^*′*^(*m*) are the genotypes of rare variant *m* of two individuals, *p*(*h*_1_, *h*_2_, *h*_3_ | *D*) and *p*(*h*_1_, *h*_2_, *h*_3_, *h*_4_ | *D*) stand for the joint probabilities of haplotypes from two individuals given *D. D* represents the available haplotypes from the same population, such as CEU in the 1000 Genomes Project. For common variants, the haplotype joint probabilities could be approximated by the product of haplotype frequencies [16]. However, this strategy is not effective for rare variants, as their MAFs are low and can be inaccurately estimated from the population. We assume the probability of the presence of a rare variant minor allele follows a Bernoulli Distribution,

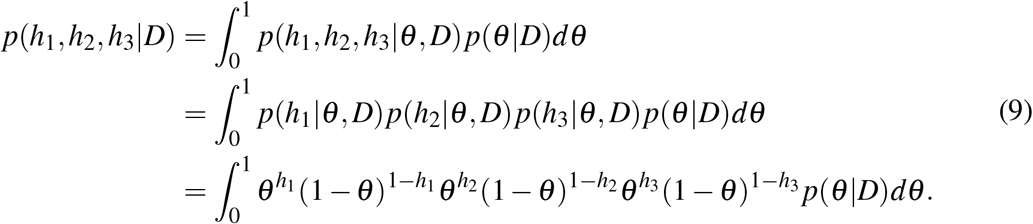

*θ* is a latent variable that estimates the parameter of the Bernoulli distribution. *p*(*θ* | *D*) follows a beta distribution and it can be expanded as

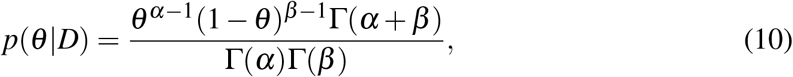

where *α* and *β* are the numbers of major and minor alleles of the reference population and test sets for the rare variant *m*. The LLR *τ*_*m*_ of the rare variant *m* is defined as

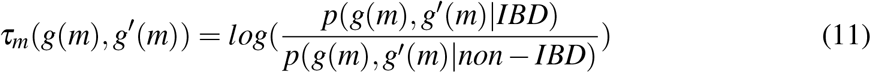

### Determination of IBD segments

IBD segments can be identified by considering the LLRs of common variants (*λ*_*b*_) and rare variants (*τ*_*m*_). If *λ*_*b*_ is larger than *T*_*LLR*_ (by default 3), the block is considered to be a long IBD segment. SILO applies a seed-and-extend algorithm to identify short IBD segments using both common variants and rare variants. In the seed-and-extend algorithm, each rare variant is assigned to a window that covers it or is the nearest to it. We define the combined-LLR *ϕ*_*w*_ of a window *w* as

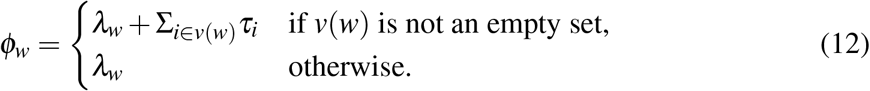

*v*(*w*) is the set of all rare variants in the window *w*. A seed is defined as a window *w* such that *ϕ*_*w*_ *>*= *T*_*LLR*_, *λ*_*w*_ *>*= *T*_*w*1_, and *w* contains rare variants with Σ_*i*_*∈*_*v*(*w*)_*τ*_*i*_ *>* 1. During the extension, SILO will first consider the left flanking window and then the right flanking window (Figure 1C). The extension procedure terminates if one of the following criteria is satisfied: (i) the summation of combined-LLRs of the involved windows becomes smaller than *T*_*LLR*_; (ii) the number of windows with *λ*_*w*_ smaller than a threshold *T*_*w*2_ exceeds the threshold 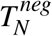, and the proportion of windows with *λ*_*w*_ smaller than *T*_*w*2_ exceeds the threshold 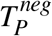; (iii) the ϕ_*w*_ is smaller than -3; (iv) the extended region is longer than 2 cM (since long IBD segments are supposed to be detected by common variants). If the extension is stopped in one direction, the extension in the opposite direction continues until it also meets one of the above criteria. The final post-extension region is regarded as a short IBD segment.

### Removal of false positive IBD segments

SILO includes a cleaning procedure to remove potential false positive IBD segments using a seed-and-extend algorithm. A seed is defined as a window *w* if *ϕ*_*w*_ *< T*_*LLR*_ and *w* contains rare variants with Σ_*i*_*∈*_*v*(*w*)_*τ*_*i*_ *<* 0. The extension procedure is similar to the one described above, but with different termination conditions. The procedure terminates if one of the following criteria is satisfied: (i) the summation of combined-LLRs of the involved windows becomes larger than or equal to *T*_*LLR*_; (ii) the *λ*_*w*_ is larger than -1; (iii) the extended region is longer than 1 cM. The overlaps of the extended regions and IBD segments are the false positive regions. SILO removes these false positive regions from the inferred IBD segments.

### Generation of simulated pedigrees

We employed ped-sim [30, 31] to simulate pedigrees on chromosome 1 (GRCh38) with some pedigree members harboring a disease susceptible variant. The founders of the simulated pedigrees were randomly selected from the unrelated white people (population code: GBR, CEU, TSI, FIN, IBS, MXL, PUR) in the 1000 Genomes Project [25, 28]. We estimated the MAF of each variant from these unrelated white people, excluding the ones selected as the founders. The disease susceptible variant was randomly selected from rare variants with MAFs below 1%. The genetic map of chromosome 1 was obtained from https://bochet.gcc.biostat.washington.edu/beagle/genetic_maps/. The male and female genetic maps were treated identical. We employed the crossover inference model using parameters provided by ped-sim for the simulation.

### Generation of simulated data with no latent IBD segments

Fastsimcoal2 [26, 27] was used to generate raw SNVs following the similar strategy used in Refined IBD [32]. In this study, we simulated a genomic region of 30 Mb. The effective population size in the current generation was 24.3 million. We sampled 1,000 individuals (2,000 haplotypes) in the current generation as the reference population, and an additional 150 individuals (300 haplotypes) in the current generation as a test set. The mutation rate was set to 2.5 × 10^−8^ and the recombination rate was 10^−8^ (1 Mb = 1 cM). There was no transition bias in simulation. We only kept common variants (MAF>= 5%) in the reference population.

To minimize the impact of latent IBD segments in the test set, which could introduce unexpected noise in performance evaluations, we adjusted the strategy used by Parente2 [16] to break up those IBD segments by dividing the genomic sequences into many small contiguous windows, each with a length of 0.05 cM. Briefly, we shuffled the haplotypes in each window of the individuals in the test set, thereby generating 150 composite individuals, where the two haplotypes of each composite individual may originate from different individuals. We assumed no IBD segments existed in these composite individuals, and they replaced the original individuals in the test set.

We selected a pair of individuals from the test set and transplanted a disease susceptible variant (MAF below 1%) along with a haplotype segment (0.1 cM to 2 cM) at a random position from one individual to the other. For the disease susceptible variant, the shared haplotype between two individuals was assigned a minor allele, while the other allele for each individual was determined based on its MAF. We also manually introduced genotype errors with a rate of 0.5% for common variants and 0.001% for the disease susceptible variant. We performed 100 simulations for each of the following IBD length intervals: 0.1 cM-0.2 cM, 0.2 cM-0.5 cM, 0.5 cM-1 cM, and 1 cM-2 cM.

### IBD inference using a real pedigree from the 1000 Genomes Project

We downloaded the haplotypes (GRCh38) of the individuals in the 1000 Genomes Project from The International Genome Sample Resource (IGSR) [25, 28]. A pedigree of Mexican descent was selected, which consisted of six members. Their relationships to one another are presented in **Figure S1** based on the pedigree information from https://www.internationalgenome.org. We also selected individuals that were unrelated and had annotated population codes of either GBR, CEU, TSI, FIN, IBS, MXL or PUR to constitute the reference population. All individuals belonging to the Mexican pedigree were removed from the reference population. As a result, we retained 674 individuals in the reference population. For quality control, we removed singletons and SNVs with missing value rates exceeding 10%, and kept only bi-allelic SNVs. In addition, we only kept common variants (MAF*≥*5%) and rare variants (MAF*≤*1%).

To validate whether these rare variants were inherited from one haplotype of a founder, NA19660 or NA19661, we applied the following criteria: (i) only one haplotype of a founder, NA19660 or NA19661, harbored a minor allele of the rare variant; (ii) NA19662, NA19685, and NA19686 all shared a minor allele in the rare variant; (iii) NA19684 harbored two major alleles in the rare variant; (iv) the corresponding haplotype segments of NA19660 (or NA19661), NA19662, NA19685, and NA19686 were identical surrounding the rare variant, with the haplotype segments containing the surrounding 40 SNVs (20 located upstream and 20 located downstream of the rare variant). Only rare variants that satisfied the criteria were selected in the subsequent analysis.

### Benchmark framework

We benchmarked the performance of SILO against four IBD detection models: hap-IBD [20], GERMLINE2 [18], TRUFFLE [22], and Parente [15]. The descriptions of these models and the specific software versions employed in our experiments are provided in **Table S4** and **Table S5**, respectively. Initially, we also included HapFABIA [33] in our benchmarking experiments; however, it was ultimately excluded from the analysis due to its inability to infer any IBD segments. This can be attributed to HapFABIA’s difficulties in classifying rare tagSNPs defined by its model when the number of individuals in the dataset is limited.

We employed default hyperparameters and tuned hyperparameters of each model for the simulated pedigres (**Table S6**-**S7**), and tuned hyperparameters for the simulated pairs with no latent IBD segments (**Table S8**-**S11**). We comparatively evaluated the performance of each model based on hit rates of disease susceptible variants (defined as as the number of variants found by an IBD detection model divided by the total number of variants), and power, precision, and *F*_1_ scores of IBD segment detection. We used the default hyperparameters of each model to identify IBD segments of the Mexican pedigree from the 1000 Genomes Project (**Table S12**), and evaluated the performance with the following metrics: (i) the potential disease susceptible variants that could be identified by a model while missed by the other models; (ii) the comparison of inferred and expected IBD segment proportions.

## Data availability

Simulated pedigrees were generated by ped-sim (https://github.com/williamslab/pedsim). Simulated pairs with no latent IBD segments were generated by fastsimcoal2 (http://cmpg.unibe.ch/software/fastsimcoal27/). The haplotypes from the 1000 Genomes Project are available at http://ftp.1000genomes.ebi.ac.uk/vol1/ftp/data_collections/1000G_2504_high_coverage/working/20220422_3202_phased_SNV_INDEL_SV/. The genetic maps of GRCh38 can be downloaded from https://bochet.gcc.biostat.washington.edu/beagle/genetic_maps/.

## Code availability

SILO’s source code and documentation are available at https://github.com/ericcombiolab/SILO.

## Acknowledgments

We would like to thank Dr. Sivan Bercovici and Prof. Serafim Batzoglou for their useful comments and supervision in the early stage of this project. We thank the Research Grants Council, Hong Kong, the Research Committee of Hong Kong Baptist University and the Interdisciplinary Research Clusters Matching Scheme for their kind support of this project. This research was partially supported by the Hong Kong Research Grant Council Early Career Scheme (HKBU 22201419), Guangdong-Hong Kong Technology Cooperation Funding Scheme (GHX/133/20SZ), HKBU Start-up Grant Tier 2 (RC-SGT2/19-20/SCI/007), HKBU IRCMS (No. IRCMS/19-20/D02) and the Guangdong Basic and Applied Basic Research Foundation (No. 2021A1515012226).

## Authors’ contributions

LZ conceived the study, CW collected data, developed the software, and conducted experiments. CW and LZ wrote the manuscript. W.P.V and L.Z. edited the manuscript. All authors reviewed the manuscript.

## Additional information

**Correspondence** and requests for materials should be addressed to Lu Zhang.

