## Supplementary materials for "Enhanced disease susceptible variant identification via short identity by descent segments"

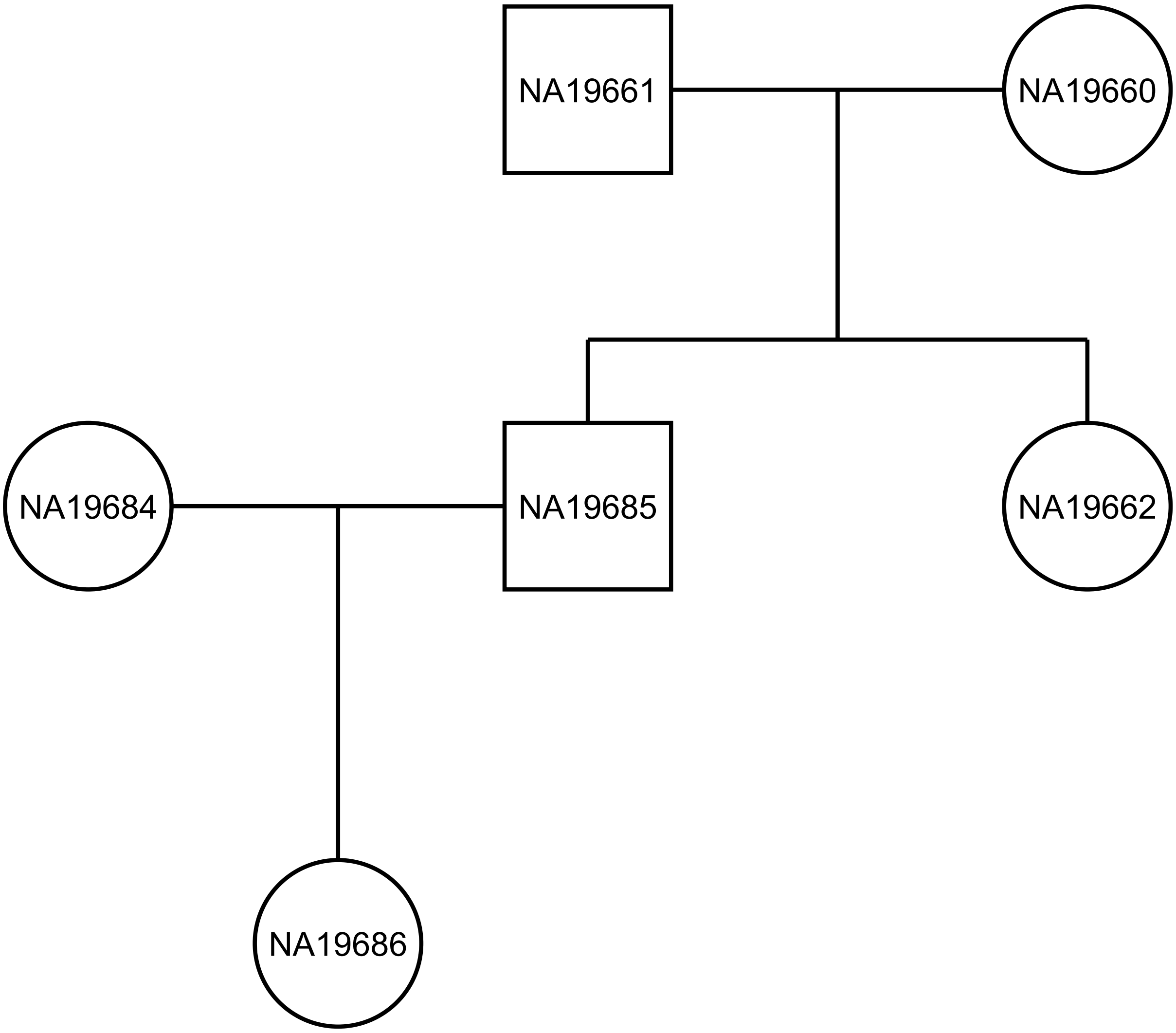


**Figure S1:** The pedigree chart of a Mexican family in the 1000 Genomes Project.

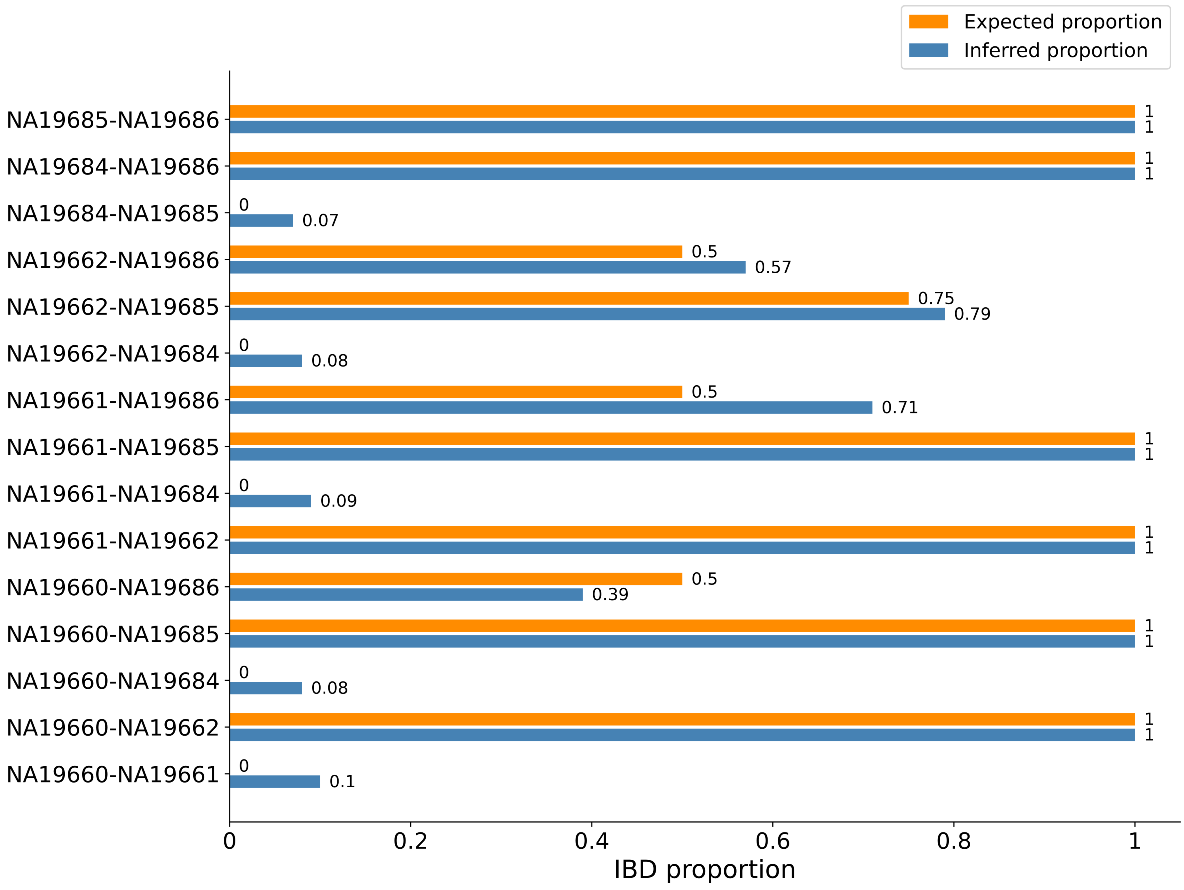


**Figure S2:** Comparison of the proportions of inferred IBD segments detected by SILO and the expected proportions of IBD segments shared among various pairs of pedigree members in the Mexican pedigree from the 1000 Genomes Project.

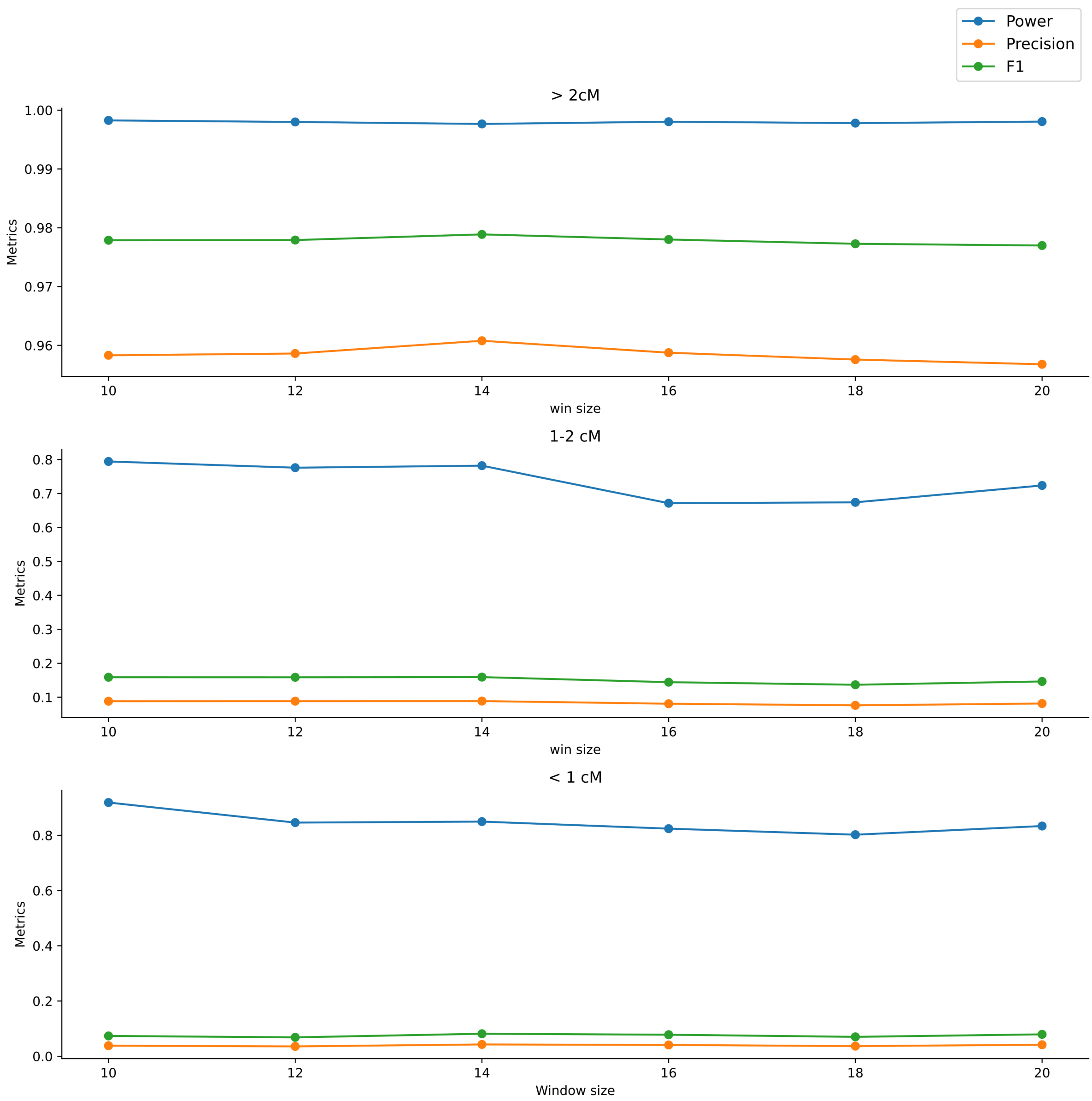


**Figure S3:** The performance of SILO with different *window size* for simulated pedigrees.

| Individual 1 | Individual 2 | SILO | Hap-IBD | GERMLINE2 | Parente | TRUFFLE |
|---|---|---|---|---|---|---|
| NA19660 | NA19661 | 0.101 | 0 | 0.022 | 0.003 | 0 |
| NA19660 | NA19662 | 0.999 | 0.833 | 0.953 | 0.997 | 0.996 |
| NA19660 | NA19684 | 0.075 | 0 | 0.013 | 0.005 | 0 |
| NA19660 | NA19685 | 0.999 | 0.853 | 0.943 | 0.997 | 1 |
| NA19660 | NA19686 | 0.389 | 0.278 | 0.326 | 0.325 | 0.308 |
| NA19661 | NA19662 | 0.999 | 0.843 | 0.944 | 0.996 | 0.989 |
| NA19661 | NA19684 | 0.090 | 0.001 | 0.016 | 0.004 | 0.001 |
| NA19661 | NA19685 | 0.999 | 0.855 | 0.943 | 0.996 | 0.997 |
| NA19661 | NA19686 | 0.710 | 0.560 | 0.637 | 0.678 | 0.673 |
| NA19662 | NA19684 | 0.079 | 0.001 | 0.020 | 0.002 | 0 |
| NA19662 | NA19685 | 0.792 | 0.655 | 0.716 | 0.746 | 0.706 |
| NA19662 | NA19686 | 0.566 | 0.42 | 0.478 | 0.491 | 0.432 |
| NA19684 | NA19685 | 0.072 | 0.001 | 0.015 | 0.008 | 0 |
| NA19684 | NA19686 | 0.999 | 0.793 | 0.952 | 0.997 | 0.997 |
| NA19685 | NA19686 | 0.999 | 0.892 | 0.949 | 0.996 | 0.998 |

**Table S1:** The mean proportions of inferred autosomal IBD segments of each model for different combination of individuals in the Mexican pedigree.

| $T_{w1}$ | $T_{w2}$ | Power | Precision | $F_1$ score |
|---|---|---|---|---|
| 0 | -0.3 | 0.737 | 0.046 | 0.087 |
| -0.1 | -0.3 | 0.840 | 0.045 | 0.085 |
| -0.2 | -0.2 | 0.793 | 0.044 | 0.083 |
| -0.2 | -0.3 | 0.850 | 0.043 | 0.082 |
| -0.2 | -0.4 | 0.880 | 0.044 | 0.084 |
| -0.3 | -0.3 | 0.879 | 0.042 | 0.080 |

**Table S2:** SILO's performance with different $T_{w1}$ and $T_{w2}$ hyperparameter settings for IBD lengths shorter than 1 cM in simulated pedigrees.

| $T_{w1}$ | $T_{w2}$ | Power | Precision | $F_1$ score |
|---|---|---|---|---|
| 0 | -0.3 | 0.578 | 0.083 | 0.145 |
| -0.1 | -0.3 | 0.702 | 0.086 | 0.153 |
| -0.2 | -0.2 | 0.635 | 0.079 | 0.141 |
| -0.2 | -0.3 | 0.782 | 0.089 | 0.160 |
| -0.2 | -0.4 | 0.707 | 0.082 | 0.147 |
| -0.3 | -0.3 | 0.721 | 0.082 | 0.147 |

**Table S3:** SILO's performance with different $T_{w1}$ and $T_{w2}$ hyperparameter settings for IBD lengths between 1 to 2 cM in simulated pedigrees.

| Method | Main algorithm | Haplotypes/Genotypes | Probabilistic model |
|---|---|---|---|
| GERMLINE2 | Hash and extension | Haplotypes | No |
| Hap-IBD | PBWT and extension | Haplotypes | No |
| HapFABIA | Biclustering | Haplotypes/Genotypes | No |
| Parente | Naive Bayes | Genotypes | Yes |
| TRUFFLE | Search long shared regions | Genotypes | No |

**Table S4:** An overview of IBD detection models. PBWT: positional Burrows-Wheeler transform.

| Method | Version or creation time |
|---|---|
| SILO | 1.0 |
| Hap-IBD | 1.0 |
| GERMLINE2 | July 2018 |
| Parente | 1.0.1 |
| TRUFFLE | 1.38 |
| HapFABIA | 1.28.0 |

**Table S5:** The versions or created time of each model.

| Method | Hyperparameter | Par1 |
|---|---|---|
| SILO | window size | 14 |
| | negative_ratio_thres | 0.15 |
| | negative_count_thres | 1 |
| | $T_{w1}$ | -0.2 |
| | $T_{w2}$ | -0.3 |
| Hap-IBD | min-seed | 2 |
| | min-output | 2 |
| | min-markers | 100 |
| GERMLINE2 | minimum match length | 1 |
| Parente | window size | 5 |
| | block length | 4 |
| TRUFFLE | length threshold | 1 |

**Table S6:** The default hyperparameters of each model in the simulated pedigrees.

| Method | Hyperparameter | Par1 |
|---|---|---|
| SILO | window size | 14 |
| | negative_ratio_thres | 0.15 |
| | negative_count_thres | 1 |
| | $T_{w1}$ | -0.2 |
| | $T_{w2}$ | -0.3 |
| Hap-IBD | min-seed | 1 |
| | min-output | 1 |
| | min-markers | 80 |
| GERMLINE2 | minimum match length | 0.4 |
| Parente | window size | 5 |
| | block length | 1.5 |
| TRUFFLE | length threshold | 0.5 |

**Table S7:** The tuned hyperparameters of each model in the simulated pedigrees.

| Method | Hyperparameter | Par1 | Par2 | Par3 | Par4 |
|---|---|---|---|---|---|
| SILO | window size | 8 | 10 | 12 | - |
| | negative_ratio_thres | $\frac{1}{10}$ | $\frac{1}{3}$ | - | - |
| | negative_count_thres | 1 | 3 | - | - |
| | $T_{w1}$ | 0 | - | - | - |
| | $T_{w2}$ | 0 | - | - | - |
| Hap-IBD | min-seed | 0.07 | 0.1 | 0.15 | 0.2 |
| | min-output | 0.07 | 0.1 | 0.15 | 0.2 |
| | min-markers | 20 | 50 | 100 | - |
| GERMLINE2 | minimum match length | 0.07 | 0.1 | 0.15 | 0.2 |
| Parente | window size | 8 | 10 | 15 | 20 |
| | block length | 0.07 | 0.1 | - | - |
| TRUFFLE | length threshold | 0.1 | 0.15 | 0.2 | - |

**Table S8:** The tuned hyperparameters of each model for IBD segments within 0.1-0.2 cM in simulated pairs with no latent IBD segments. The best one (highest $F_1$ scores) listed above for each model was selected for performance evaluation.

| Method | Hyperparameter | Par1 | Par2 | Par3 | Par4 | Par5 | Par6 |
|---|---|---|---|---|---|---|---|
| SILO | window size | 8 | 10 | 12 | - | - | - |
| | negative_ratio_thres | $\frac{1}{10}$ | $\frac{1}{3}$ | - | - | - | - |
| | negative_count_thres | 1 | 3 | - | - | - | - |
| | $T_{w1}$ | 0 | - | - | - | - | - |
| | $T_{w2}$ | 0 | - | - | - | - | - |
| Hap-IBD | min-seed | 0.1 | 0.15 | 0.2 | 0.3 | 0.4 | 0.5 |
| | min-output | 0.1 | 0.15 | 0.2 | 0.3 | 0.4 | 0.5 |
| | min-markers | 20 | 50 | 100 | - | - | - |
| GERMLINE2 | minimum match length | 0.1 | 0.15 | 0.2 | 0.3 | 0.4 | 0.5 |
| Parente | window size | 8 | 10 | 15 | 20 | - | - |
| | block length | 0.2 | 0.3 | - | - | - | - |
| TRUFFLE | length threshold | 0.2 | 0.3 | 0.4 | 0.5 | - | - |

**Table S9:** The tuned hyperparameters of each model for IBD segments within 0.2-0.5 cM in simulated pairs with no latent IBD segments. The best one (highest $F_1$ scores) listed above for each model was selected for performance evaluation.

| Method | Hyperparameter | Par1 | Par2 | Par3 | Par4 | Par5 | Par6 | Par7 | Par8 |
|---|---|---|---|---|---|---|---|---|---|
| SILO | window size | 8 | 10 | 12 | - | - | - | - | - |
| | negative_ratio_thres | $\frac{1}{10}$ | $\frac{1}{3}$ | - | - | - | - | - | - |
| | negative_count_thres | 1 | 3 | - | - | - | - | - | - |
| | $T_{w1}$ | 0 | - | - | - | - | - | - | - |
| | $T_{w2}$ | 0 | - | - | - | - | - | - | - |
| Hap-IBD | min-seed | 0.3 | 0.4 | 0.5 | 0.6 | 0.7 | 0.8 | 0.9 | 1 |
| | min-output | 0.3 | 0.4 | 0.5 | 0.6 | 0.7 | 0.8 | 0.9 | 1 |
| | min-markers | 20 | 50 | 100 | - | - | - | - | - |
| GERMLINE2 | minimum match length | 0.3 | 0.4 | 0.5 | 0.6 | 0.7 | 0.8 | 0.9 | 1 |
| Parente | window size | 8 | 10 | 15 | 20 | - | - | - | - |
| | block length | 0.5 | 0.6 | - | - | - | - | - | - |
| TRUFFLE | length threshold | 0.3 | 0.4 | 0.5 | 0.6 | 0.7 | 0.8 | 0.9 | 1 |

**Table S10:** The tuned hyperparameters of each model for IBD segments within 0.5-1 cM in simulated pairs with no latent IBD segments. The best one (highest $F_1$ scores) listed above for each model was selected for performance evaluation.

| Method | Hyperparameter | Par1 | Par2 | Par3 | Par4 | Par5 | Par6 | Par7 | Par8 |
|---|---|---|---|---|---|---|---|---|---|
| SILO | window size | 8 | 10 | 12 | - | - | - | - | - |
| | negative_ratio_thres | $\frac{1}{10}$ | $\frac{1}{3}$ | - | - | - | - | - | - |
| | negative_count_thres | 1 | 3 | - | - | - | - | - | - |
| | $T_{w1}$ | 0 | - | - | - | - | - | - | - |
| | $T_{w2}$ | 0 | - | - | - | - | - | - | - |
| Hap-IBD | min-seed | 0.3 | 0.5 | 0.8 | 1 | 1.3 | 1.5 | 1.8 | 2 |
| | min-output | 0.3 | 0.5 | 0.8 | 1 | 1.3 | 1.5 | 1.8 | 2 |
| | min-markers | 20 | 50 | 100 | - | - | - | - | - |
| GERMLINE2 | minimum match length | 0.3 | 0.5 | 0.8 | 1 | 1.3 | 1.5 | 1.8 | 2 |
| Parente | window size | 8 | 10 | 15 | 20 | - | - | - | - |
| | block length | 1 | 1.3 | - | - | - | - | - | - |
| TRUFFLE | length threshold | 0.3 | 0.5 | 0.8 | 1 | 1.3 | 1.5 | 1.8 | 2 |

**Table S11:** The tuned hyperparameters of each model for IBD segments within 1-2 cM in simulated pairs with no latent IBD segments. The best one (highest $F_1$ scores) listed above for each model was selected for performance evaluation.

| Method | Hyperparameter | Par1 |
|---|---|---|
| SILO | window size | 14 |
| | negative_ratio_thres | 0.15 |
| | negative_count_thres | 1 |
| | $T_{w1}$ | -0.2 |
| | $T_{w2}$ | -0.3 |
| Hap-IBD | min-seed | 2 |
| | min-output | 2 |
| | min-markers | 100 |
| GERMLINE2 | minimum match length | 1 |
| Parente | window size | 5 |
| | block length | 4 |
| TRUFFLE | length threshold | 1 |

**Table S12:** The hyperparameter settings of each model in the 1000 Genomes Project.
